# Unique and conserved endoplasmic reticulum stress responses in neuroendocrine cells

**DOI:** 10.1101/2025.08.29.672971

**Authors:** Karina Rodrigues-dos-Santos, Gitanjali Roy, Anna Geisinger, Sahiti Somalraju, Travis S. Johnson, Michael A. Kalwat

## Abstract

Endocrine cells are dedicated to the production and processing of hormones, from peptides to small molecules, to regulate key physiological processes, including glucose homeostasis and metabolism. Because of this relatively high productivity, endo-crine cells must handle a variety of stresses from oxidative stress to the unfolded protein response of the endoplasmic reticulum (UPR^ER^). While much is known about the major pathways regulating the UPR^ER^, the roles of endocrine cell type-specific, context-dependent, and time-dependent transcriptional changes are not well explored. To identify unique and shared responses to the UPR^ER^ across a subset of endocrine cell types, we tested representative lines for β-cells (insulin), α-cells (glucagon), δ-cells (somatostatin), X/A-cells (ghrelin), L-cells (glucagon-like peptide 1 (GLP1)), and thyrotropes (thyroid hormone and thyroglobulin). We exposed each cell type to the canonical ER stressor thapsigargin for 6 and 24 h, or vehicle for 24 h and performed mRNA sequencing. Analysis of the data showed all lines responded to thapsigargin. Comparisons of differentially expressed genes between each line revealed both shared and unique transcriptional signatures. These data represent a valuable mineable set of candidate genes that may have cell type-specific functions during the UPR^ER^ and have the potential to lead to a new understanding of how different endocrine cells mitigate or succumb to ER stress.

## 1. Introduction

The unfolded protein response of the endoplasmic reticulum (UPR^ER^) is conserved across eukaryota and is engaged when the burden of secretory protein production ex-ceeds capacity, inducing a gene program to mitigate stress or, if mitigation is not achieved, induce apoptosis [1, 2]. The UPR^ER^ has been an attractive source of pharmaco logical targets for diseases, including diabetes and cancer [3, 4]. Dedicated secretory cell types, like neuroendocrine cells, have a unique reliance on these stress pathways and represent useful models for studying the mechanisms underlying ER stress and the subsequent activation of UPR^ER^ [5-7]. The UPR^ER^ involves activation of three canonical signaling pathways: PERK-eIF2α-ATF4; IRE1α-XBP1s; and ATF6[8]. The UPR^ER^ is a double-edged sword in certain endocrine cells like pancreatic islet β-cells. β-cells require the UPR^ER^ for normal function, but in diseases like type 1 diabetes (T1D) and type 2 diabetes (T2D), chronic engagement of the UPR^ER^ may exacerbate β-cell failure. The relative cell type-specific signatures of UPR^ER^ are not well-defined across disparate types of endocrine cells. Characterizing these unique signature genes may enable the identification of novel cell-type-specific biomarkers and targets, as well as new insights into the underlying biology, thereby improving the understanding of the UPR^ER^ in general.

In this study, we have generated a transcriptomic resource comparing the responses of a panel of dedicated secretory cell types to the canonical ER stressor, thapsigargin. We performed computational analyses to cross-compare all cell types and identify unique and shared transcriptomic responses. We have focused here on the unique UPR^ER^ responses of β-cells and α-cells, given the preponderance of data indicating α-cells have a distinct response to stress, which may contribute to their survival in T1D [9]. However, we expect that these data will be useful across multiple fields and have provided an interactive web tool to facilitate UPR^ER^ comparisons.

## 2. Materials and Methods

### Compounds

Thapsigargin was from Adipogen (AGCN20003M001). All other chemicals were from reputable sources including Sigma-Aldrich and Fisher Scientific.

### Cell culture, treatments, and RNA isolation

MIN6 β-cells (RRID: CVCL_0431) were cultured in high glucose DMEM containing 10% fetal bovine serum (FBS), 50 µM β-mercaptoethanol, 4 mM L-glutamine, 1 mM pyruvate, 100 U/mL penicillin, and 100 µg/mL streptomycin. The mouse intestinal L cell line GLUTag (RRID: CVCL_J406) was cultured in low glucose DMEM containing 10% FBS, 4 mM L-glutamine, 1 mM pyruvate, 100 U/mL penicillin, and 100 µg/mL streptomycin. Human somatostatinoma QGP-1 (RRID: CVCL_3143) cells [10] were cultured in RPMI-1640 with 10% FBS, 1 mM pyruvate, 4 mM L-glutamine, 100 U/mL penicillin, and 100 µg/mL streptomycin. The mouse ghrelinoma cell line MGN3 (RRID: CVCL_C4PL) [11] was cultured in high glucose DMEM supplemented with 10% FBS and containing 4 mM L-glutamine, 100 U/ml penicillin and 100 µg/ml streptomycin. Tissue culture plates were pre-coated with diluted Matrigel (1:20) for 1 h at 37°C prior to seeding. The rat thyroid cell line PCCL3 (RRID: CVCL_6712) [6] was cultured in Coons F-12 supplemented with 1 mIU/mL thyrotropin, 1 µg/mL insulin, 5 µg/mL apo-transferrin, 1 nM hydrocortisone, 5% FBS, 100 U/mL penicillin, and 100 µg/mL streptomycin. The mouse pancreatic alpha cell line αTC1 clone 6 (αTC1-6) (RRID: CVCL_B036) was previously described [12]. αTC1-6 cells were cultured in 15 mM glucose DMEM supplemented with 15 mM HEPES, 0.1 mM non-essential amino acids, 10% FBS, 0.03% BSA, 100 U/mL penicillin, and 100 µg/mL streptomycin. GLUTag, QGP-1, PCCL3, aTC1, MGN3 and MIN6 cells were plated in 6-well dishes and exposed to DMSO 0.1% or thapsigargin 100 nM (GLUTag, QGP1, MGN3 and MIN6) or 500 nM (PCCL3 and αTC1-6) for 6 and 24 hours before harvesting for RNA purification, downstream gene expression analysis and RNAseq. A higher dose of thapsigargin was used for PCCL3 and αTC1-6 cells based in part on the observed cellular morphology after 24 h of treatment and on published data showing effects of 100 nM thapsigargin requiring 48 h exposure[13] to substantially impact PCCL3 and that α-cells are known to be resistant to ER stress conditions [9].

### Human islet culture and treatment

Cadaveric human islets were obtained through the Integrated Islet Distribution Program (IIDP) and Prodo Labs. Islets were isolated by the affiliated islet isolation center and cultured in PIM medium (PIM-R001GMP, Prodo Labs) supplemented with glutamine/glutathione (PIM-G001GMP, Prodo Labs), 5% Human AB serum (100512, Gemini Bio Products), and ciprofloxacin (61-277RG, Cellgro, Inc) at 37°C and 5% CO_2_ until shipping at 4°C overnight. Human islets were cultured upon receipt in complete CMRL-1066 (containing 1 g/L (5.5 mM) glucose, 10% FBS, 100 U/ml penicillin, 100 µg/ml streptomycin, 292 µg/mL L-glutamine). Human islet information and donor metadata are provided in Table S1. For drug treatments, ∼50 human islets were hand-picked under a dissection microscope, transferred to low-binding 1.5 ml tubes, and cultured in 500 µL of complete CMRL-1066 medium containing DMSO (0.1%) for 24 h or thapsigargin (1µM) for 6 and 24 h.

### RNA isolation and RT-qPCR validation

ER stress was induced in cell lines and human islets using thapsigargin as described for 6 and 24 h. After indicated treatments in cell lines, medium was removed from cells, and lysis buffer (with β-mercaptoethanol) was added to cells. Cells were scraped, and lysates were transferred to 1.5 mL tubes on ice and then transferred to −80°C for storage and RNA was later isolated using the Aurum Total RNA Mini kit (Bio-Rad). For human islets, islets were harvested and RNA isolated using the Quick-RNA Microprep (Zymo). RNA concentration was measured using a Nanodrop spectrophotometer and verified to have A260/280 ratios > 2.0. RNA (1µg for cell lines, 500ng for islets) was converted into complementary DNA (cDNA) using the iScript cDNA synthesis kit (Bio-Rad) following manufacturer instructions. cDNAs were diluted 10-fold, and 1 µL was used per qPCR reaction to confirm the induction of ER stress responses. One microliter of diluted cDNA was used in 10 µL quantitative polymerase chain reactions using 2X SYBR Bio-Rad master mix and 250 nM of each primer. Reactions were run in 384-well format on QuantStudio 5 (Thermo). qPCR data were analyzed using the QuantStudio software with *18S* RNA as the reference gene for cell lines and *ACTB* and *VAPA* as reference genes for human islets [14]. Relative expression was calculated by the 2-ΔΔCt method. All primer sequences are provided in **Table S2**.

### Transcriptomics

300ng of RNA was submitted to the Indiana University Center for Medical Genomics for mRNA-seq. Total RNA samples were first evaluated for their quantity and quality using Agilent Bioanalyzer 2100. All samples had good quality with RIN (RNA Integrity Number) ≥9. 100 nanograms of total RNA was used for library preparation with the KAPA mRNA Hyperprep Kit (KK8581) on Biomek following the manufacturer’s protocol. Each uniquely dual-indexed library was quantified and quality assessed by Qubit and Agilent TapeStation, and multiple libraries were pooled in equal molarity. The pooled libraries were sequenced with 2X 100bp paired-end configuration on an Illumina NovaSeq 6000 sequencer using the v1.5 reagent kit. Samples had an average read depth of ∼52.7 million reads/sample. The sequencing reads were first quality checked using FastQC (v.0.11.5, Babraham Bioinformatics, Cambridge, UK) for quality control. The sequence data were then mapped to either the mouse reference genome mm10, the human reference genome hg38, or the rat reference genome rn6 using the RNA-seq aligner STAR (v.2.710a) [15] with the following parameter: “--outSAMmapqUnique 60”. To evaluate the quality of the RNA-seq data, the number of reads that fell into different annotated regions (exonic, intronic, splicing junction, intergenic, promoter, UTR, etc.) of the reference genome was assessed using bam-stats (from NGSUtilsJ v.0.4.17) [16]. Uniquely mapped reads were used to quantify the gene level expression employing featureCounts (subread v.2.0.3) [17] with the following parameters: “-s 2 -p –countReadPairs Q 10”. Transcripts per million (TPM) were calculated using length values determined by using the “makeTxDbFromGFF” and “exonsBy” functions in the “GenomicFeatures” library and the “reduce” function in the “GenomicRanges” library in R to find the length of the union of non-overlapping exons for each gene [18].

### Data processing and analysis

edgeR 4.4.2 was used for the differential expression analysis of the RNA-seq data [19]. Cutoffs were set for FDR < 0.05 and fold change (|log_2_fold-change| > 1) to determine which genes were up-regulated or down-regulated. Pairwise comparisons were performed between the control group (DMSO), thapsigargin 6 h, and thapsigargin 24 h groups, generating a list of differentially expressed genes (DEGs) for each time point and cell line, which were used to create volcano plots in R using ggplot2 v3.5.2. For comparisons of DEGs across cell lines, we generated upset plots using the UpSetR package [20]. Genes changed uniquely in each cell line were extracted and these lists were analyzed in R using Gene Ontology and Gene Set Enrichment Analysis (fGSEA in R, GO terms string-db [21] and EnrichR [22]). R scripts are numbered in the order they were used in the workflow and are provided on GitHub (https://github.com/kalwatlab/endocrine-ER-stress/).

### Interactive web tool design

To generate a web tool to interact with the RNAseq data, we used shiny in R to integrate the data and analyses from our pipeline. The app includes tabs for volcano plot, gene set dot plot generation, data table, pairwise cell line comparisons, interactive upset plots, and GSEA results. The web tool is available at https://diabetes-detectives.shinyapps.io/endocrine-ER-stress/.

### Statistical Analysis

Graphed data are expressed as mean ± SD. Data were evaluated using one- or two-way ANOVA as indicated with appropriate post-hoc tests and considered significant if P < 0.05. GraphPad Prism 10 was used to perform statistical tests for qPCR graphs. Statistics for transcriptomics data was performed in R using edgeR with the design being set to model.matrix(∼0 + group, data = dge_list$samples) and fit with the glmQLFit() function.

## 3. Results

### 3.1 Thapsigargin induces ER stress in multiple endocrine cell lines from distinct tissue types

To study endocrine cell type-specific responses to ER stress, we selected cell lines that represent the pancreatic islet (α-cells: αTC1-6; β-cells: MIN6; δ-cells: QGP1), the gut (L-cells: GLUTag; X/A-cells: MGN3-1), and the thyroid (thyrocytes: PCCL3) (Fig 1A). To induce the UPR^ER^ in different endocrine cell types, we used a canonical ER stressor, thapsigargin, which inhibits the SERCA2 Ca^2+^ ATPase, causing ER Ca^2+^ depletion. We confirmed by RT-qPCR that treatment with thapsigargin for either 6 or 24h induced canonical UPR^ER^ genes, including *Hspa5* (BiP) and *Ddit3* (CHOP), across all cell lines (Fig 1B-G). Thapsigargin also significantly reduced expression of cell-type-specific hormones in some cell lines, including *Ins1/2* in MIN6 β-cells, *Gcg* in αTC1 α-cells, and thyroglobulin in PCCL3 thyrocytes (Fig 1B,C,G).

**Figure 1.**
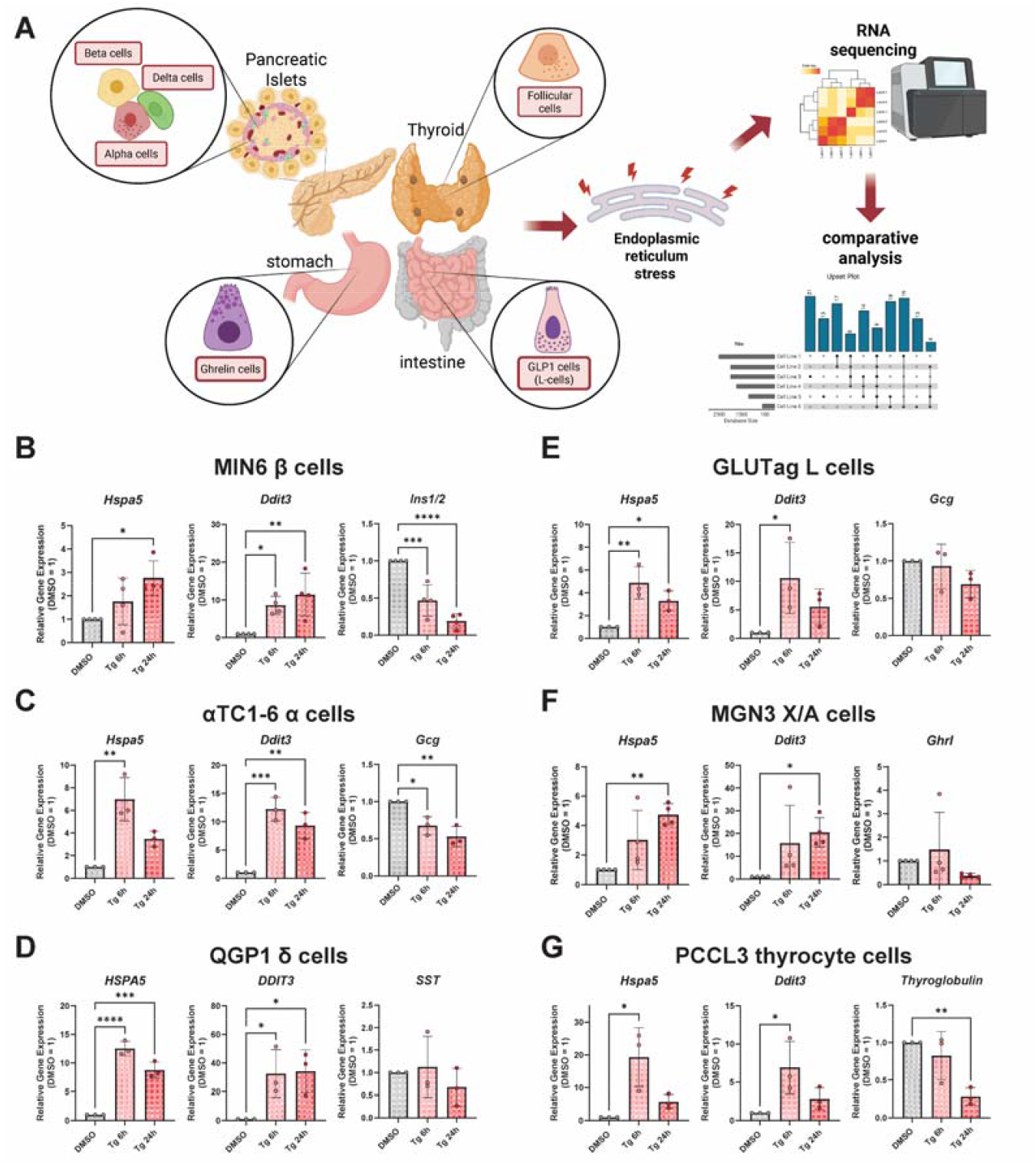
Selection of representative endocrine cell lines and validation of induction of UPR^ER^ genes. **A)** Cartoon model depicting selection of endocrine cell types and their tissues of origin. Figure created in Biorender. **B-G**) qPCR of genes in the lines for validation. Data are the mean ± SD of N = 3-4. ^*^, P<0.05; ^**^, P<0.01; ^***^,P<0.001 vs DMSO by one-way ANOVA with Dunnett’s multiple comparisons test.

Having validated that Tg induces the UPR^ER^ in each of the selected lines, we performed bulk RNA-seq on all samples (DMSO, Tg 6h, Tg 24h; N=3 per cell line). The transcriptomic data were processed and analyzed separately for each cell line using edgeR to identify differentially-expressed genes for Tg 6h vs DMSO and Tg 24h vs DMSO for each cell line (**Fig 2A-F**). All edgeR results were merged into a supplemental table (**Table S3**). Endocrine cell lines expressed their expected cell type-specific markers [23-26], including *Gcg, Mafb*, and *Arx* for α-cells; *Gcg, Chga*, and *Nts* for L-cells; *Ghrl, Acsl1, Cartpt*, and *Prg4* for X/A (ghrelin) cells; *Ins1, Mafa, Slc2a2* for β-cells; and *Tg* (thyroglobulin), *Tpo*, and *Tshr* for thyrocytes; and *Sst, Hhex*, and *Bche* for δ-cells (**Fig 2G-H**). At both 6 and 24h exposure to thapsigargin, these cell type markers were the most significantly decreased by Tg in α-cells, β-cell and thyrocytes, while ghrelin cell, L-cell and δ-cell markers were less affected (**Fig 2G-H**).

**Figure 2.**
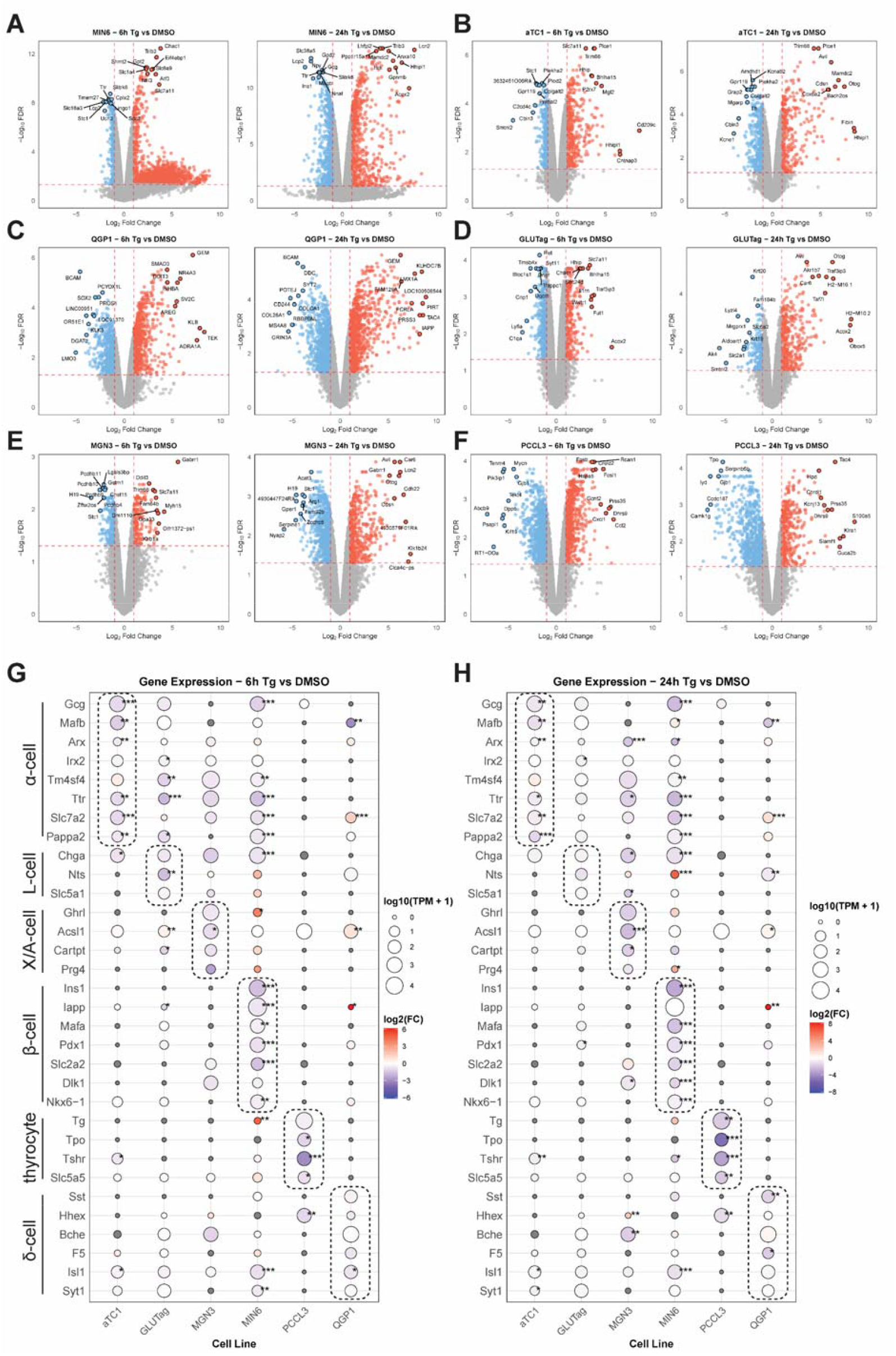
Endocrine cell type responses to thapsigargin-induced ER stress. Volcano plots of differentially-expressed genes (DEGs) at 6 and 24h of thapsigargin (Tg) treatment for **A**) MIN6 β-cells, **B**) αTC1 α-cells, **C**) QGP1 δ-cells, **D**) GLUTag L-cells, **E**) MGN3 X/A-cells, and **F**) PCCL3 thyrocytes. Cell type-specific marker gene expression for each cell line is shown for 6 h (**G**) and 24 h (**H**) of Tg treatment. Data are the mean of three independent biological replicates. ^*^, FDR<0.05; ^**^, FDR<0.01; ^***^,FDR<0.001 vs DMSO by edgeR analysis.

We also confirmed that, as has been shown previously [27, 28], Tg induced the UPR^ER^ in human islets, which are comprised mainly of β-, α-, and δ-cells. We assessed this by comparing the expression of ER stress genes at the 6 and 24 h time points in our cell lines (**Fig S2A**) to that of human islets treated similarly (**Fig. S2B**). Following these validations, we next used multiple approaches to examine the similarities and differences in responses to Tg across the different endocrine lines.

### 3.2 Integrating cell-type-specific UPR^ER^ transcriptomic responses identifies common and unique gene signatures

To find distinct and shared gene sets, we used an upset plot analysis to identify the number of DEGs unique to each cell line, as well as those in common between all comparisons of cell lines (**Fig 3A, Table S4**). Genes that are regulated by Tg in common across all tested cell lines included known factors like *Ddit3* (encoding CHOP), *Ppp1r15a, Slc7a11* (**Fig 3B**). As expected, this common gene set was enriched for ontology terms related to ER protein processing and response to stress **(Fig 3C**). We also extracted cell line-unique DEGs (**Table S5**). A subset of the top up- and down-regulated unique genes from the Tg 24 h time point for β-cells and α-cells is shown (**Fig 3D, E**). For the MIN6 β-cell-unique genes, the molecular function GO terms included tRNA synthetase activity, RNA modifications like N6-methyladenosine, and solute/ion transport (**Fig S2C, Table S6**). In contrast, for αTC1 cell-unique genes, the molecular function GO terms included carboxypeptidase activity and aldehyde dehydrogenase activity (**Fig S2D, Table S6**).

**Figure 3.**
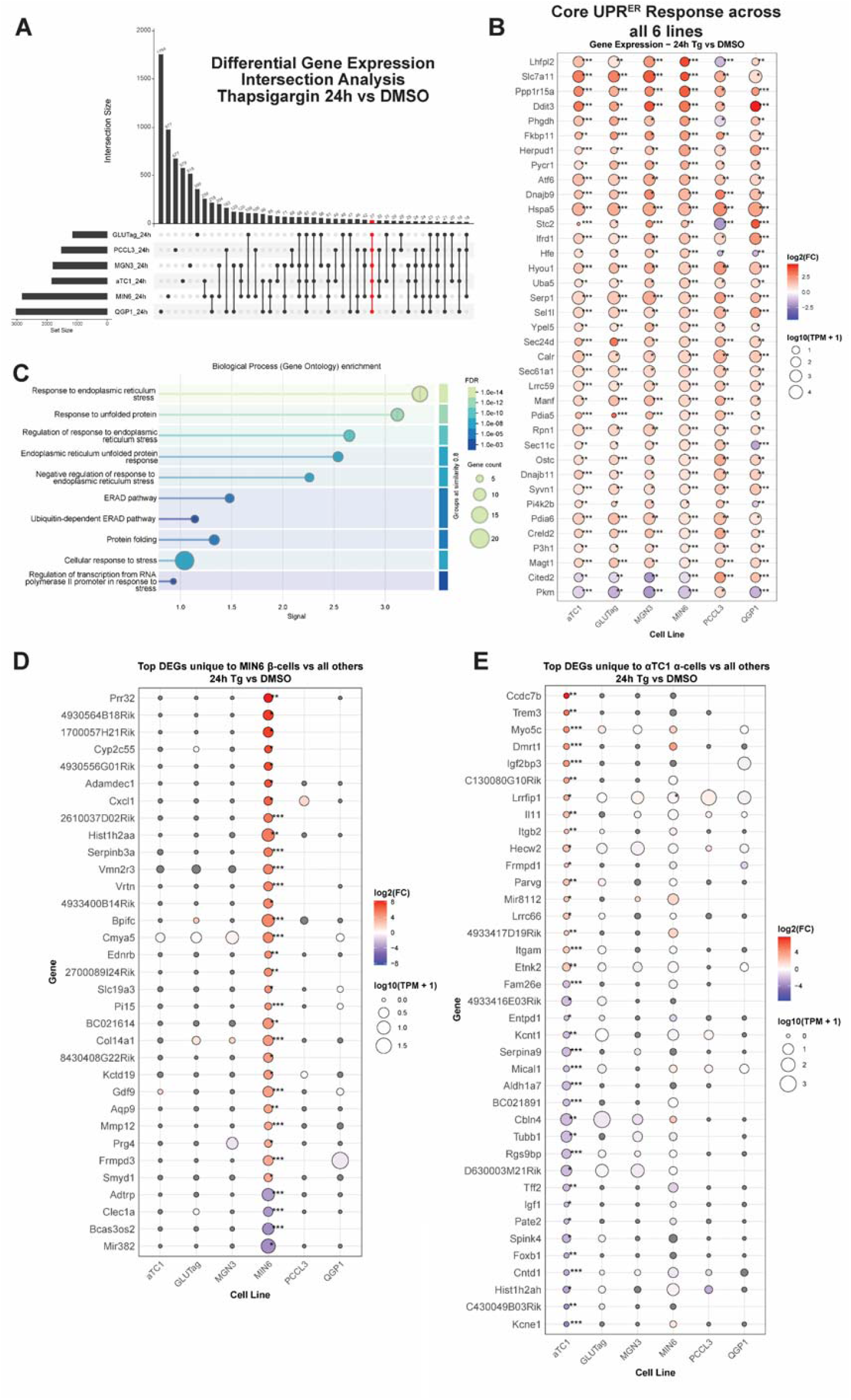
**A**) Upset plot analysis comparing significantly differentially expressed genes (DEGs_ after 24h of Tg treatment across all 6 cell types. Highlighted in red is the set of DEGs that is common to all cell lines. **B**) Dot plot of DEGs which are shared among all lines. **C**) Gene ontology term enrichment from String-DB of the gene set shown in (**B**). **D**) Subset of the top DEGs that are uniquely changed in MIN6 cells at Tg 24h. **E**) Subset of the top DEGs that are uniquely changed in αTC1 cells at Tg 24h. In all dot plots, dot size is proportional to average expression level in transcripts per million (TPMs) and are colored by log_2_FC. ^*^, FDR<0.05; ^**^, FDR<0.01; ^***^,FDR<0.001 vs DMSO by edgeR analysis.

### 3.3 Cell type comparisons identify concordant and discordant responses to UPR^ER^ in β-cells versus α-cells

To identify concordant and discordant gene expression, we used pairwise log_2_FC comparisons. We correlated the pairwise log_2_FC of DEGs for all lines (**Fig S3**). The analyses are browsable and exportable via our web app. As an example, we focus on the comparison between MIN6 β-cells and αTC1-6 α-cells (**Fig 4A and S4A, Table S7**). We found that while many genes altered by thapsigargin were concordant between these two cell types, there was a subset of discordant genes (**Fig 4B-C**). One of the most discordant genes that was upregulated in α-cells, but downregulated in β-cells, was *Atp2a3* which encodes sarcoplasmic/endoplasmic reticulum Ca^2+^ ATPase 3 (SERCA3) (**Fig 4B**). The SERCA proteins are the main target of thapsigargin, which can inhibit all SERCA isoforms [29]. Interestingly, among all endocrine cells tested, αTC1 and GLUTag cells were the only lines which significantly induced SERCA3. On the other hand, one of the most discordant genes upregulated in β-cells but downregulated in α-cells was *Sprr1a* (**Fig 4C**). Additionally, when considering the 6 and 24 h time points together, several discordantly upregulated genes in β-cells had to do with cargo receptor activity, including *Cubn, Dmbt1, Tmprss3, Megf10*, and *Loxl3* (**Fig 4C and S4C**), suggesting β-cells alter their intracellular cargo handing or trafficking substantially differently than α-cells.

**Figure 4.**
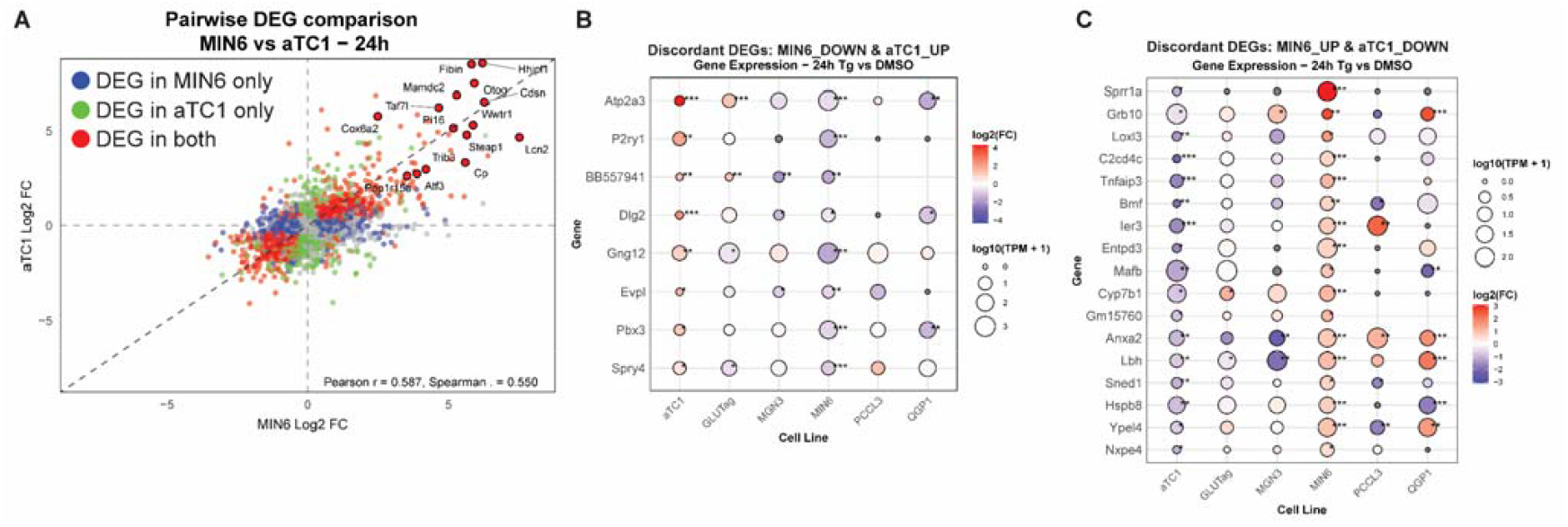
**A**) Pairwise comparison of log_2_FC of all genes changed by Tg 24 h in MIN6 β-cells versus αTC1 α-cells. Differentially expressed genes (DEGs) in MIN6 only are in blue, αTC1 only are in green, and DEG in both are in red. **B**) Discordant DEGs that are downregulated in MIN6 but upregulated in αTC1 after 24 h of Tg treatment. **C**) Discordant DEGs that are upregulated in MIN6 but downregulated in αTC1 after 24 h of Tg treatment. FDR<0.05; ^**^, FDR<0.01; ^***^,FDR<0.001 vs DMSO by edgeR analysis.

Finally, we compared our β-cell and α-cell data with a recent single-cell RNAseq dataset from Maestas, et al., where human islets were treated with different stressors including Tg, bafilomycin A, and cytokines for 48 h [30]. The authors identified 6 genes that were commonly upregulated in α-, β-, and δ-cells: *CIB1, ERP44, HSP90B1, NEAT1, SELK*, and *VMP1*. We see general agreement in our dataset with these genes being upregulated in αTC1, MIN6, and QGP-1 (**Fig S4D**). One exception is the lncRNA NEAT1 which was downregulated in αTC1.

### 3.4 Generation of an interactive transcriptomics browser for endocrine UPR^ER^

To provide a resource to the UPR^ER^ field, we have integrated our RNAseq data into an interactive web tool using shiny [31], accessible at https://diabetes-detectives.shinyapps.io/endocrine-ER-stress/. This tool has functions for browsing volcano plots (**Fig 5A**), pairwise comparisons of log_2_FC data between cell lines (**Fig 5B**), a custom gene set dot plot generation tool (**Fig 5C**), GSEA results across cell lines (**Fig 5D**), a filterable data table viewer for all cell line edgeR results and TPMs, and an interactive upset plot viewer (**Fig 5**).

**Figure 5.**
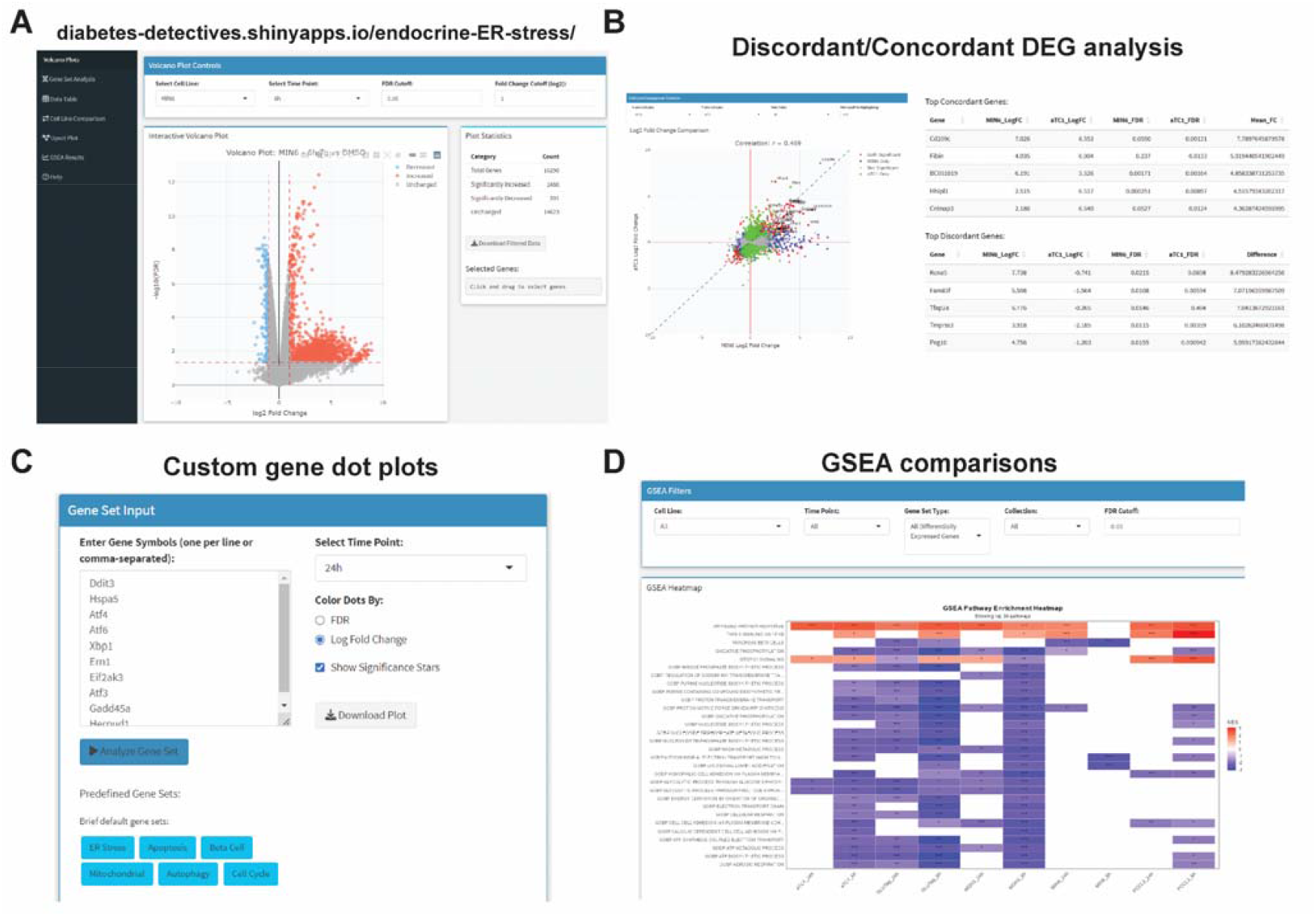
Interactive transcriptomics explorer for endocrine ER stress data. **A**) The data explorer has multiple interactive plot types (e.g. volcano, pairwise log2FC, upset plots). **B**) Pairwise comparisons allow for export of plots and of concordant/discordant genes for selected cell line and time point. **C**) Custom gene set analysis that generates dot plots. Genes are kept in the order given, size of dot is log_10_(TPMs+1) and colored by log_2_FC. **D**) Filterable GSEA results for each time point and cell type with export capabilities.

## 4. Discussion

The UPR^ER^ is engaged in T1D and T2D in the β-cell and is an attractive source of pharmacological targets for diseases, including diabetes [4, 32, 33]. Compared to related cell types, β-cells have a unique reliance upon the UPR^ER^ as they handle the daily task of insulin production [34]. The differences and similarities of β-cells from other specialized secretory cells is in part due to their requirement for exquisite balance in this pathway. Heterogeneity of genetic and environmental backgrounds likely influences the resistance or susceptibility of β-cells to stress. For example, β-cells require XBP1 to maintain identity [35]; however, IRE1α deletion can help protect β-cells in T1D models [36]. The β-cell UPR^ER^ is also engaged in monogenic forms of diabetes, such as mutant insulin-induced diabetes of youth (MIDY)[37], where mutations in the INS gene cause misfolding of the insulin protein, leading to activation of UPR^ER^ and β-cell dysfunction [38]. Therefore, investigations into the effects of pharmacological or genetic modulation of the UPR^ER^ in β-cells has broad mechanistic and therapeutic implications.

Pancreatic islet β-cells and α-cells are well known to differ in their responses to stress. For example, in T1D the immune system recognizes β-cells and destroys them, but does not target α-cells [39]. The contributors to this phenomenon have been speculated to include higher expression of anti-apoptotic, viral recognition, and innate immune response genes in α-cells, as well as a relatively higher response to ER stress in β-cells [9]. In our study, we have identified candidate genes that may contribute to α-cell-specific vs β-cell-specific responses to canonical ER stress. For example, we found that *Atp2a3* (SERCA3) is induced in α-cells but downregulated in β-cells (Fig 4B). SERCA3 was previously shown to be expressed in mouse pancreatic islet β-cells, but absent in islet α-cells [40], although human islet scRNAseq suggests that α-cells and δ-cells express the *ATP2A3* gene [25, 41]. Perhaps α-cell up-regulation of SERCA3 could contribute to their known resistance to ER stress [9]. Indeed, small molecule SERCA activators are under investigation as potential β-cell therapeutics in diabetes [42, 43]. Another gene selectively upregulated in α-cells was *Igf2bp3* (Fig 3E). Igf2bp3 plays a role in binding to N6-methyladenosine (m6A) modified mRNA (ref). m6A modifications have become of increasing interest in T1D and T2D biology due to their potential regulatory role in pancreatic β-cells [44, 45]. The role(s) for m6A regulation in α-cells and how *Igf2bp3* may be important during islet stress are unknown.

One of the most discordant genes upregulated in the β-cell, but downregulated in α-cells, was *Sprr1a*, which encodes small proline-rich protein 1A. SPRR1A is involved in keratinocyte function as a structural protein [46], although new functions have recently been identified. In mice with autophagy-deficient β-cells (inducible knockout of *Atg7* only in β-cells), Sprr1a was found to be one of the most highly upregulated gene after 2-6 weeks of induced knockout [47]. The authors confirmed this at the protein level in islets. Sprr1a was also upregulated in db/db mouse islets and in wild-type mice subjected to chemically-induced insulin resistance. Taken together, these findings support that *Sprr1a* is a stress-responsive gene, possibly specific to β-cells. None of the other endocrine cell types we tested had increased *Sprr1a* expression.

To our knowledge, this study is the first comparative transcriptomics analysis of multiple distinct endocrine cell type responses in an ER stress model at multiple time points. These data represent a useful resource for multiple fields, including general secretory cell biology, neuroendocrine tumors (NETs), and metabolic and endocrine diseases.

### Limitations of the study

Our raw data is from cell lines, although we compared the data to primary human islet gene expression data. Induction of ER stress in our study depended on a pharmacological agent, thapsigargin. While this compound has been used for decades for this purpose, it may not fully recapitulate the UPR^ER^ that occurs normally *in vivo*. Additionally, inferences from this data are also dependent on RNA expression, which does not always reflect protein expression. Future work will require validation studies for these signature genes at the protein level in specific endocrine cell types from primary tissues, and using additional ER stress model systems.

## Supplementary Materials

The following supporting information can be downloaded at: www.mdpi.com/xxx/s1, Figure S1: title; Table S1: title; Video S1: title.

## Author Contributions

Conceptualization, K.R.S. and M.A.K.; methodology, K.R.S. and M.A.K.; software, K.R.S., S.S., T.S.J., and M.A.K.; formal analysis, K.R.S. and M.A.K.; investigation, K.R.S., G.R., and M.A.K.; resources, A.G.; data curation, K.R.S. and M.A.K.; writing—original draft preparation, K.R.S. and M.A.K.; writing—review and editing, K.R.S., G.R., and M.A.K.; visualization, K.R.S. and M.A.K.; supervision, M.A.K, funding acquisition, M.A.K. All authors have read and agreed to the published version of the manuscript.

## Funding

This research and the APC were funded in part by a Pilot & Feasibility grant administered by the Indiana University Center for Diabetes and Metabolic Diseases (P30 DK097512) and internal support from the IBRI. The Center for Medical Genomics at Indiana University School of Medicine is partially supported by the Indiana University Grand Challenges Precision Health Initiative.

## Institutional Review Board Statement

Not applicable.

## Data Availability Statement

RNA-seq data associated with this manuscript is publicly available under NCBI GEO identifier GSE238017. Code is provided on Github (link). All other supplementary materials are provided as part of this manuscript or its online repository.

## Acknowledgments

PCCL3 cells were provided by Dr. Peter Arvan (University of Michigan). QGP1 cells were provided by Dr. Dawn E. Quelle (University of Iowa). MGN3-1 cells were provided by Dr. Hiroshi Iwakura (Wakayama Medical University, Japan). GLUTag L-cell line was created by Dr. Dan Drucker (Mount Sinai Hospital, Toronto CA) and was provided by the lab of George G. Holz (Upstate Medical University). αTC1-6 cells were provided by Dr. Michael Roth (UT Southwestern). MIN6 β-cells were a gift from Dr. John Hutton. Sequencing analysis was conducted at the Center for Medical Genomics at Indiana University School of Medicine, which is partially supported by the Indiana University Grand Challenges Precision Health Initiative. We thank personnel in both the Center for Computational Biology and Bioinformatics and the Center for Medical Genomics, directed by Dr. Yunlong Liu.

## Conflicts of Interest

The authors declare no conflicts of interest. The funders had no role in the design of the study; in the collection, analyses, or interpretation of data; in the writing of the manuscript; or in the decision to publish the results.

## Abbreviations

The: following abbreviations are used in this manuscript
UPR^ER^: Unfolded protein response of the endoplasmic reticulum
T1D: Type 1 diabetes
T2D: Type 2 diabetes
GSEA: Gene set enrichment analysis
GLP1: Glucagon-like peptide 1

## Disclaimer/Publisher’s Note

The statements, opinions and data contained in all publications are solely those of the individual author(s) and contributor(s) and not of MDPI and/or the editor(s). MDPI and/or the editor(s) disclaim responsibility for any injury to people or property resulting from any ideas, methods, instructions or products referred to in the content.

## Notes

https://github.com/kalwatlab/endocrine-ER-stress

https://diabetes-detectives.shinyapps.io/endocrine-ER-stress/

### Competing Interest Statement

The authors have declared no competing interest.

