## Supplementary Figures for "Unique and conserved endoplasmic reticulum stress responses in neuroendocrine cells"

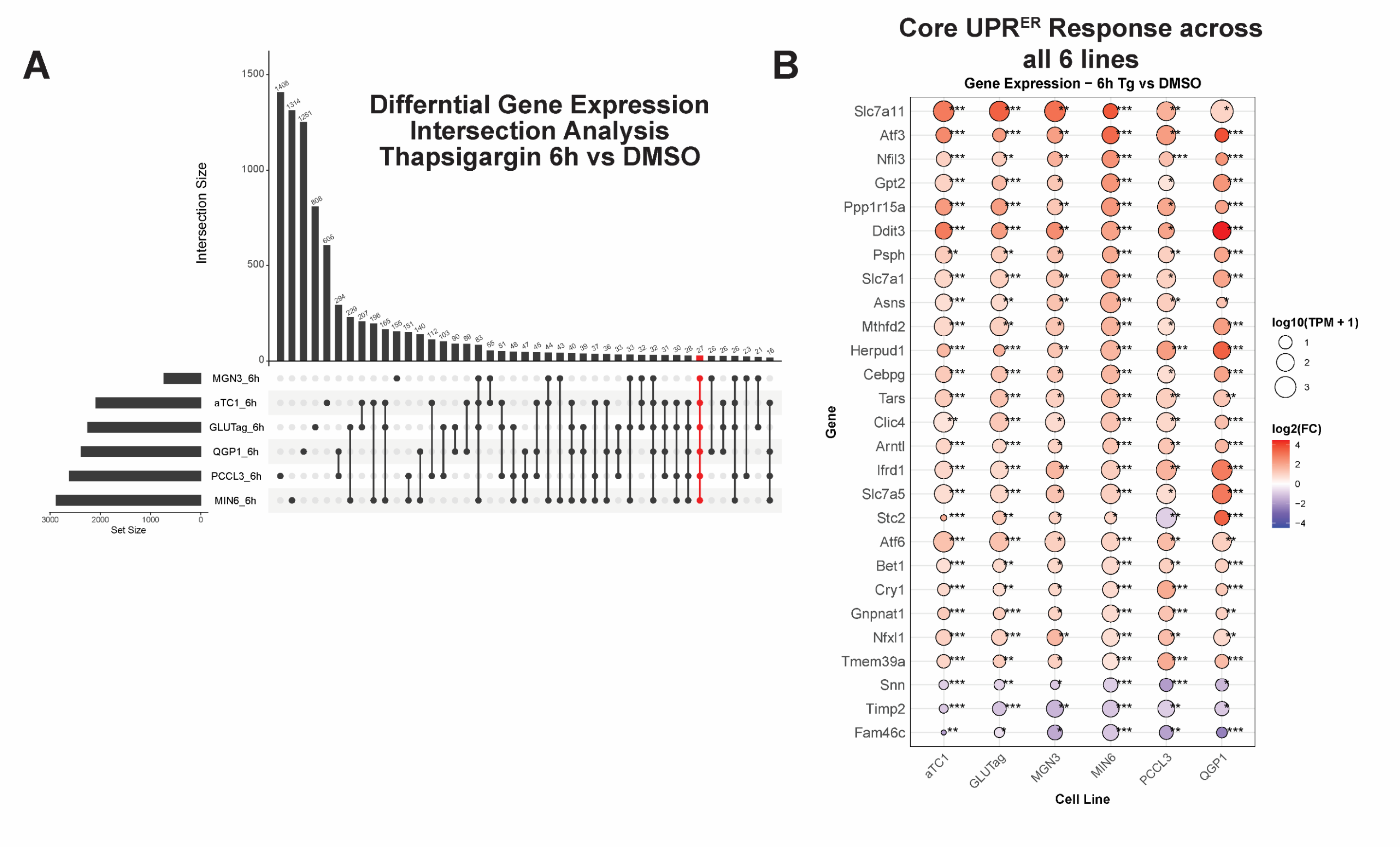


**Figure S1 – Related to Figure 3. A)** Upset plot of DEGs in all cell lines at Tg 6h. Highlighted in red is the set of DEGs that is common to all cell lines. **B)** Dot plot of DEGs shared between all 6 cell lines at Tg 6h. FDR<0.05; **, FDR<0.01; ***,FDR<0.001 vs DMSO by edgeR analysis.

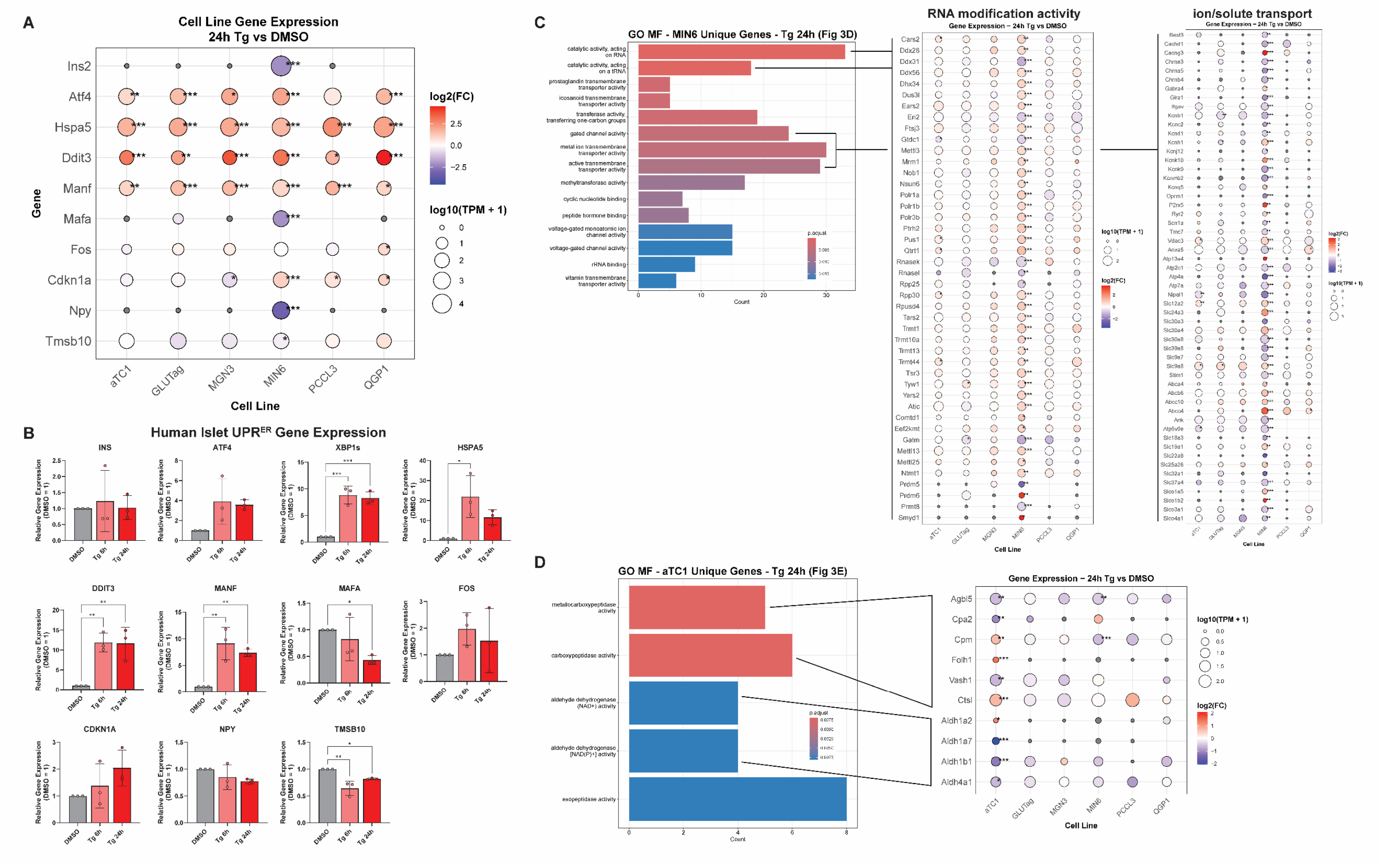


**Figure S2 – Related to Figure 3. A)** Subset of ER stress and β-cell genes for comparison to gene expression in human islets treated with Tg for 6 and 24h in (**B**). **B)** Human islets treated with Tg (1μM) for 6 or 24 h and gene expression quantified by RT-qPCR. *, P< 0.05 vs DMSO by one-way ANOVA with Dunnett's multiple comparisons test. **C)** Molecular Function (MF) Gene Ontology (GO) enrichment for entire list of MIN6 unique DEGs (related to Fig 3D) at Tg 24 h. Dot plots show gene expression for genes within selected terms for RNA modifying activity and ion/solute transport activity. **D)** Similar to (**C**), MF GO enrichment terms for αTC1 unique DEGs at Tg 24 h. Dot plots show expression for genes comprising the carboxypeptidase activity terms and aldehyde dehydrogenase activity terms. FDR<0.05; **, FDR<0.01; ***,FDR<0.001 vs DMSO by edgeR analysis.

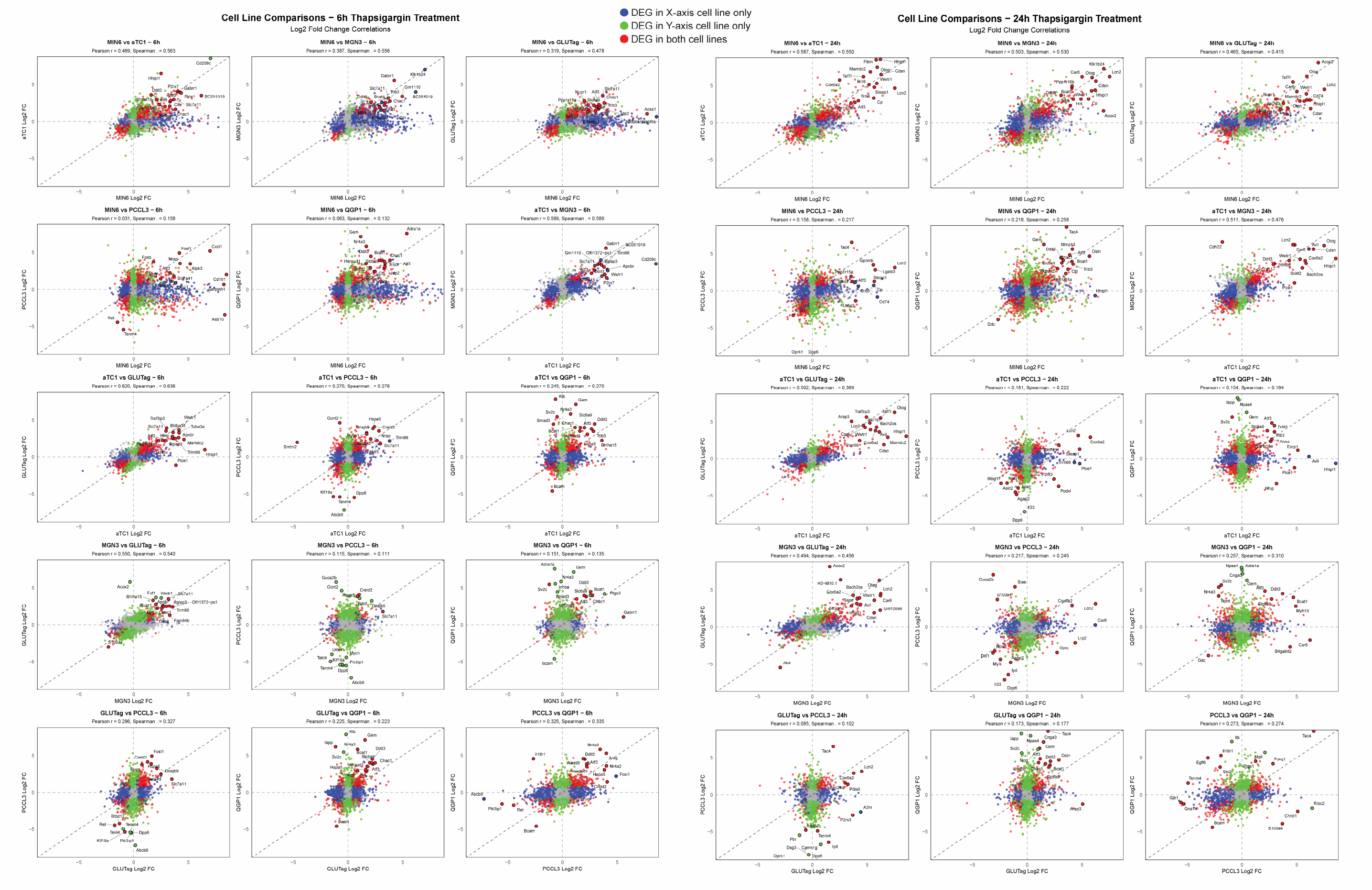


**Figure S3 – related to Figure 4.** Pairwise comparison of log2FC gene expression data for every combination of the 6 tested cell lines. Tg 6h data shown on the left and Tg 24 h data shown on the right. Differentially expressed genes (DEGs) in the X-axis cell line only are in blue, the Y-axis cell line only are in green, and red color is used for genes that are DEGs in both lines.

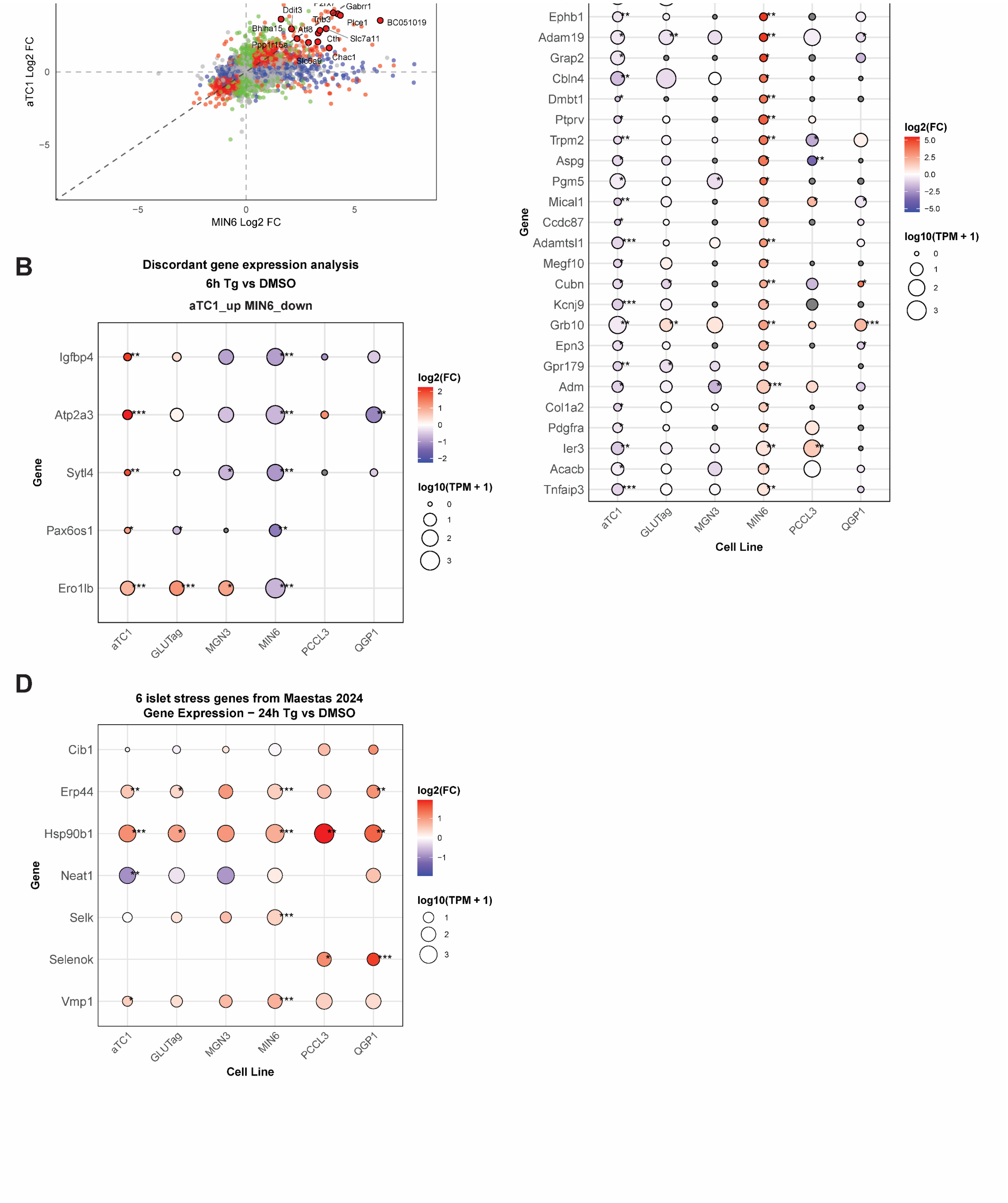


**Figure S4 – related to Figure 4. A)** Larger plot for pairwise comparison of MIN6 and αTC1 cells treated with Tg for 6 h, as seen in Fig S3. **B)** Discordant DEGs that are downregulated in MIN6 but upregulated in αTC1 after 6 h of Tg treatment. **C)** Discordant DEGs that are upregulated in MIN6 but downregulated in αTC1 after 6 h of Tg treatment. **D)** Maestas, et al. previously identified 6 core stress genes that were upregulated in human islet α-, β-, and δ-cells. The plot shows expression of these genes at Tg 24h for each of our cell lines. Selk is also known as Selenok depending the database, so both names are shown to capture the expression data. FDR<0.05; **, FDR<0.01; ***,FDR<0.001 vs DMSO by edgeR analysis.
