## Supplementary material for "Unique and conserved endoplasmic reticulum stress responses in neuroendocrine cells": Table S1

**humanChecklist for reporting human islet preparations used in research**

Adapted from Hart NJ, Powers AC (2018) Progress, challenges, and suggestions for using human islets to understand islet biology and human diabetes. Diabetologia <https://doi.org/10.1007/s00125-018-4772-2>

| **Islet preparation** | **1** | **2** | **3** | **4** | **5** | **6** | **7** | **8^a^** |
| --- | --- | --- | --- | --- | --- | --- | --- | --- |
| **MANDATORY INFORMATION** | | | | | | | | |
| Unique identifier | HP-23153-01 | HP-23166-01 | SAMN34033792 |  |  |  |  |  |
| Donor age (years) | 51 | 58 | 39 |  |  |  |  |  |
| Donor sex (M/F) | Female | Female | Male |  |  |  |  |  |
| Donor BMI (kg/m^2^) | 26.4 | 28.8 | 33.4 |  |  |  |  |  |
| Donor HbA_1c_ or other measure of blood glucose control | 5.3% | 5.6% | 5.0% |  |  |  |  |  |
| Origin/source of islets^b^ |  |  |  |  |  |  |  |  |
| Islet isolation centre | Prodo Labs | Prodo Labs | Southern California Islet Cell Resource Center |  |  |  |  |  |
| Donor history of diabetes? Please select yes/no from drop down list | No | No | No |  |  |  |  |  |
| **If Yes, complete the next two lines if this information is available** | | | | | | | | |
| Diabetes duration (years) |  |  |  |  |  |  |  |  |
| Glucose-lowering therapy at time of death^c^ |  |  |  |  |  |  |  |  |
| **RECOMMENDED INFORMATION** | | | | | | | | |
| Donor cause of death | stroke | stroke | head trauma |  |  |  |  |  |
| Warm ischaemia time (h) |  |  |  |  |  |  |  |  |
| Cold ischaemia time (h) |  |  | 12h 41m |  |  |  |  |  |
| Estimated purity (%) | 85 | 80-85 | 80 |  |  |  |  |  |
| Estimated viability (%) | 95 | 95 | 97 |  |  |  |  |  |
| Total culture time (h)^d^ |  |  |  |  |  |  |  |  |
| Glucose-stimulated insulin secretion (static culture by IIDP)^e^ |  |  | 2.2 |  |  |  |  |  |
| Handpicked to purity? Please select yes/no from drop down list | Yes | Yes | Yes |  |  |  |  |  |
| Additional notes:  Related to Figure: |  |  |  |  |  |  |  |  |
| Additional notes  -Culture time prior to shipment |  |  | 2d 7h |  |  |  |  |  |

^a^If you have used more than eight islet preparations, please complete additional forms as necessary

^b^For example, IIDP, ECIT, Alberta IsletCore

^c^Please specify the therapy/therapies

^d^Time of islet culture at the isolation centre, during shipment and at the receiving laboratory

^e^Please specify the test and the results
